# Insect homolog of oxytocin/vasopressin associated with parenting of males but not females in a subsocial beetle

**DOI:** 10.1101/2022.07.06.499045

**Authors:** Ahva L. Potticary, Christopher B. Cunningham, Elizabeth C. McKinney, Patricia J. Moore, Amsale T. Belay, Allen J. Moore

## Abstract

Parental care is thought to evolve through modification of behavioral precursors, which predicts that the mechanistic changes occur in the genes underlying those traits. The duplicated gene system of oxytocin/vasopressin has been broadly co-opted across vertebrates to influence parenting, from a pre-duplication ancestral role in water balance. It remains unclear whether co-option of these genes for parenting is limited to vertebrates. Here, we experimentally tested for associations between *inotocin* gene expression and water balance, parental acceptance of offspring, and active parenting in the subsocial beetle *Nicrophorus orbicollis*, to test whether a single copy homologue, *inotocin*, has similarly been co-opted for parental care in a species with elaborate parenting. As expected, *inotocin* was associated with water balance in both sexes. *Inotocin* expression increased around sexual maturation in both males and females, although more clearly in males. Finally, we found that expression of *inotocin* was not associated with acceptance of larvae but was associated with a transition to male but not female parenting. Moreover, level of offspring provisioning behavior and gene expression were positively correlated in males but uncorrelated in females. Our results suggest a broad co-option of this system for parenting that may have existed prior to gene duplication, and that inotocin may be associated with flexibility in parenting behavior.

**Impact Summary:** Oxytocin/vasopressin are amongst the most studied neuropeptides in vertebrates, influencing water balance, mating interactions, and most notably, social bonding. This gene pair evolved from a duplication in the vertebrate lineage of an ancestral vasopressin-like gene. Are the multiple social effects in vertebrates due to this duplication, or are social influences also ancestral? Here, we demonstrate that, in a biparental social beetle with a single copy, inotocin is associated with social interactions between fathers and offspring as well as being associated with the ancestral role of water balance in both males and females. In vertebrates, both oxytocin and vasopressin have been shown to impact social interactions in both sexes, although often showing sex-specificity in their action within species. Our results suggest that this system may have been co-opted for parenting prior to gene duplication and may facilitate flexibility in caring behavior.

## Introduction

Complex behaviors like parental care evolve through the co-option of existing behaviors and their underlying genetic mechanisms (Tallamy 1984; Cunningham et al. 2017; Benowitz et al. 2018; Moore and Benowitz 2019). However, gene networks are often robust to changes in individual genes, such that multiple genetic mechanisms may produce the same phenotypes (Manceau et al. 2010; Berens et al. 2015; Barghi et al. 2019; Cunningham et al. 2021; Berendzen et al. 2022). Thus, it is unclear whether the same genes are co-opted for parental care across taxa. This is particularly the case with the neuropeptides oxytocin and vasopressin. While water balance is an ancestral function of this system (Fujino et al. 1999; Ukena et al. 2008; Koto et al. 2019), both neuropeptides have been co-opted across vertebrate taxa to influence multiple functions and are best known for their influence on parental care and especially direct parent-offspring interactions like offspring feeding (Fig. 1, SI Table 1). These effects are often sex-specific (Fig. 1; SI Table 1) and can show remarkable complementarity in their activity within and across species (Stoop 2012). Given the diversity of phenotypic effects (Fig. 1), sex-specificity, and complementarity of action of oxytocin and vasopressin, the gene duplication in vertebrates may have facilitated this diversity of gene function in social behaviors like parental care. Yet, most taxa possess only a single homologue and show similar parental care behaviors in both sexes. How ubiquitous is the co-option of these genes for complex parental care behaviors? Is the sex-specificity maintained across taxa? A test of these questions can be made using an invertebrate with parental care that has a single copy homologue of oxytocin/vasopressin.

**Figure 1.**
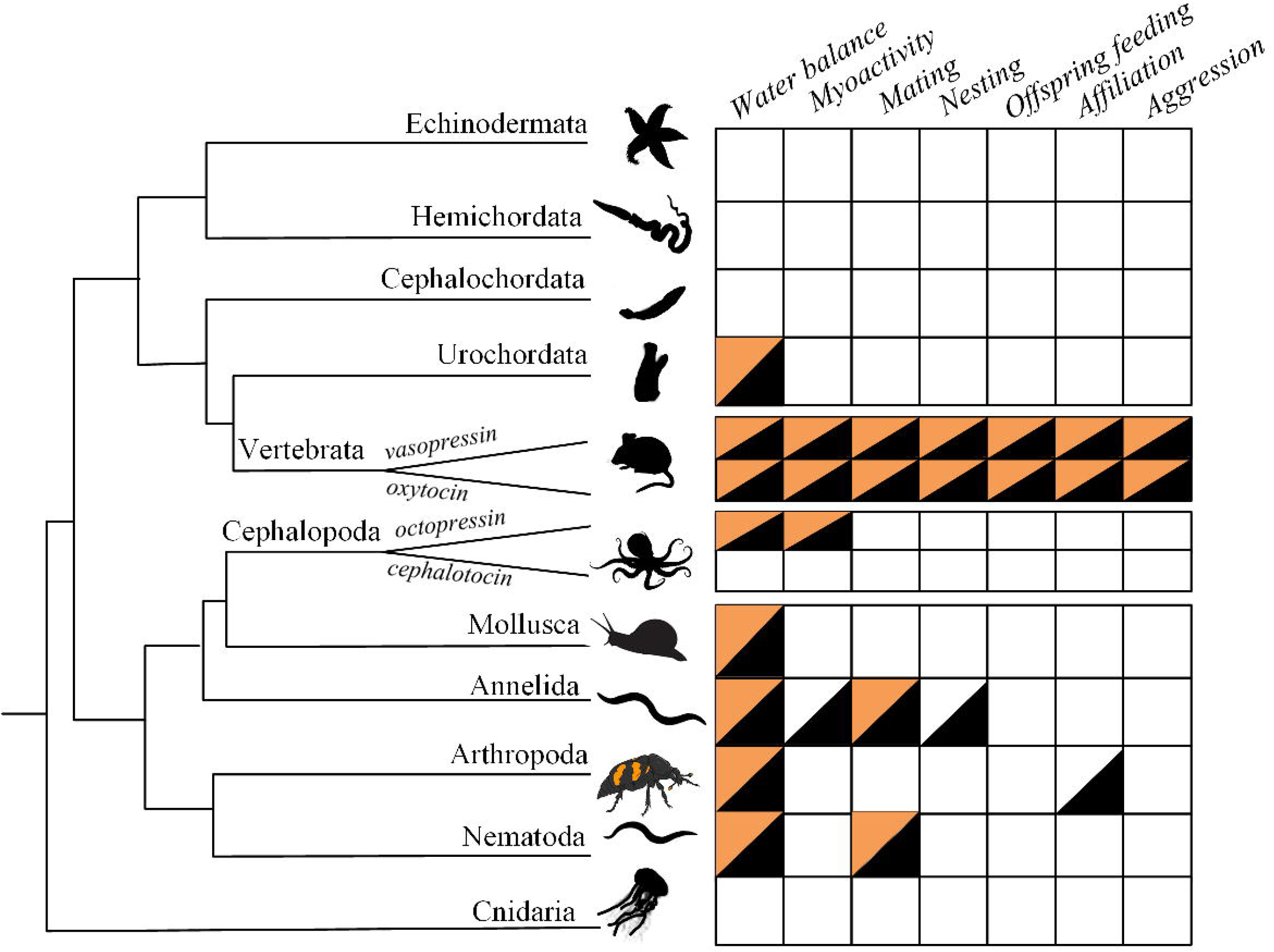
Oxytocin/vasopressin family evolved from duplication event. An ancestral vasopressin-like neuropeptide was duplicated in the common ancestor of jawed vertebrates, and another duplication occurred in the cephalopods. Duplication events denoted by labelled branches. Neuropeptides of the oxytocin/vasopressin group have been found across wide taxonomic groups (those represented on phylogeny) although the functions of these molecules are not characterized for many taxa. Functions associated with these neuropeptides include water balance, myoactivity (i.e., muscle activity, such as in egg laying), nesting, offspring care, affiliation, and aggression. Where known, functions for these homologues are indicated next to the taxonomic group they function in. Black triangles represent an association between that neuropeptide and function in females, while orange triangles indicate functionality in males. Note that the branch lengths in the tree are arbitrary, as well as the marked timing of the duplication events in vertebrata and cephaloda. Phylogeny image modified from (Elphick et al. 2018). References supporting each designation are included in Supplementary Table 1.

A single copy homologue - inotocin – exists in insects (Aikins et al. 2008; Stafflinger et al. 2008). While inotocin has been broadly associated with some social contexts in insects (Liutkevičiūtė et al. 2018; Muratspahić et al. 2020; Fetter-Pruneda et al. 2021; Nagy et al. 2021), it is unclear whether it has been specifically co-opted to influence complex parental care behaviors as most insects do not show elaborate care and prolonged parent-offspring associations as in vertebrates. Burying beetles in the genus *Nicrophorus* are an exception; parents not only have extended associations with their offspring abut also feed their offspring by regurgitating digested carrion to begging young (Scott 1998). Both males and females can care for their offspring as single parents or together. Thus, burying beetles provide a unique opportunity to determine whether inotocin is associated with similar patterns of parental care as observed in vertebrates (Fig. 1), and an evolutionary co-option of these neuropeptides for care. Here, we focus on how *inotocin* (*IT*) and *inotocin receptor* (*ITR*) expression are associated with parenting in the biparental burying beetle *Nicrophorus orbicollis*.

All burying beetles require small animal carcasses to breed, and all exhibit extensive elaborate parental care (Scott 1998). Upon finding a carcass, parents bury it, remove the exterior (e.g., fur), and coat it with an exudate to prevent decomposition. Females lay eggs in the soil and upon hatching, larvae crawl to the carcass. Burying beetles have temporal kin recognition; parents accept and feed larvae only after a certain amount of time has elapsed since egg laying has occurred (Oldekop et al. 2007). If larvae appear on the carcass too early, parents do not provide care to the offspring and eat them instead. Parental care in burying beetles involves both direct (feeding of regurgitated food to their offspring) and indirect parental care (defense and maintenance of carcass). In *N. orbicollis*, biparental or uniparental beetles raise larvae equally effectively (Benowitz and Moore 2016). Larvae of *N. orbicollis* cannot survive without parental care and under biparental conditions, females are highly consistent in their parenting, while males show more environmentally dependent care behavior (e.g., loss of partner; Benowitz and Moore 2016; Moss and Moore 2021). Parental behavior in *N. orbicollis* thus requires egg laying, a transition from infanticide to acceptance of larvae, nesting behavior, and direct larval feeding, concordant with the behavioral functions of oxytocin/vasopressin in vertebrates (Fig. 1).

We leveraged variation across the reproductive cycle of *N. orbicollis* to address whether *IT* and *ITR* gene expression in the head are associated with parenting as well as other functions of the oxytocin/vasopressin in vertebrates (including water balance). As expected, *IT* and *ITR* were related to water balance in both males and females. Expression also changed with sexual development in both sexes. Unexpectedly, variation in *IT* expression was associated with variation in male but not female parental care. Together, our results suggest that co-option of the oxytocin/vasopressin system for parenting is taxonomically widespread and may have existed pre-duplication.

## Material and Methods

### Insect Colony and Husbandry

The *Nicrophorus orbicollis* we used originated from an outbred colony regularly supplemented with wild-caught beetles from Whitehall Forest, Athens, GA, USA. We maintain this colony in incubators at 22 ± 1°C on a 14:10 hour light:dark cycle to simulate Georgia summer breeding conditions. All individuals used were of known parentage, age, and rearing conditions. We housed beetles individually at larval dispersal in 9-cm-diameter by 4-cm-deep circular compostable deli containers (Eco-Products) filled with ~2.5 cm of commercial soil. Once we placed larvae into individual containers, they had no further social interactions until they were collected for the age series or allocated into the behavioral or physiological treatments. We defined individuals as age Day 1 on the day of eclosion. We fed adult beetles organic ground beef ad libitum twice a week.

### Inotocin and Water Balance

We experimentally manipulated humidity to determine whether changes in *IT* or *ITR* in the head may be associated with water balance. We measured the relative humidity of 20 pairs of breeding *N. orbicollis* (10 at the nest/egg stage and 10 at the larval stage) that had not been opened for one week to capture the range of humidity and temperature that occurs over the breeding cycle on a decomposing mouse. We used a handheld Preciva Humidity and Temperature Reader (HT154001) to determine the relative humidity range. Low Humidity treatments were maintained between 8-25 % (15.39 ± 1.3 % RH) and High Humidity conditions were maintained between 80-95% (84.8 ± 1.3 % RH). On Day 16, we transferred beetles to an individual container with perforations nested inside of a breeding box (breeding boxes used throughout experiments were plastic containers; Pioneer Plastics, Dixon, KY) sealed with parafilm. Individuals in the Low Humidity experiment were kept without soil to ensure that they could not mitigate exposure by burrowing into the soil, while individuals in the High Humidity treatment were contained in an individual container with approximately two cm of potting soil. We simulated Low and High humidity conditions by charging the outside box with either a) 12.5 g of CaSO4 (Drierite, Hammond Drierite Co., Xena, OH) for the Low Humidity treatment, or b) a 3 x 4” cotton strip soaked to saturation with deionized water for the High Humidity. We left beetles in Low Humidity or High Humidity treatments for 8 hours to provide sufficient time for water stress to occur yet ensure that beetles survived until the end of the treatment period (Bedick et al. 2006). We collected heads of n = 10 individuals of each sex for each humidity treatment.

### Inotocin and Social Behavior

#### Development

We collected whole heads early post-adult eclosion (Day 2-3), pre-sexual maturity (Day 6-7), after sexual maturity and early reproductive age (Day 15-16), prime reproductive age (Day 21-23), late reproductive age (Day 30-31), and late-stage (Day 35-36) to test the hypothesis that *IT* changes over development to enable a shift to more social life stages. This age series corresponds to the timing when beetles were collected for each behavior state and the water balance experiment (SI Fig. 1). We collected n = 10 individuals of each sex for all ages.

#### Accept and Infanticide

We placed a mated pair (14-15 days post-eclosion) into a plastic breeding box filled with approximately two cm of potting soil moistened with distilled water and containing a freshly thawed mouse (19-25 g, Rodent Pro, Inglefield, IN, U.S.A.). Breeding boxes were returned to the incubator and kept on a darkened shelf, as this species breeds underground. We checked breeding boxes for eggs every 24 hours, and once eggs were observed (first batch), we moved the focal female and the brood ball to a new breeding box filled with approximately two cm of potting soil. The focal male was kept separate from the female and brood ball for 24 hours and then we returned the male to the breeding box with the female. Once the male and female were reunited, breeding boxes were again checked daily for new eggs (second batch).

We tested the hypothesis that *IT* influences the transition from aggression to affiliation towards larvae by manipulating the temporal recognition system of *Nicrophorus* (Oldekop et al. 2007; Benowitz et al. 2015), as parents will eat larvae that arrive before the usual embryonic period is complete (~ 96 hours in *N. orbicollis*). A pair was moved to a new box following egg laying (see above) to ensure that larvae did not arrive naturally at the carcass. We then introduced larvae to the focal pair either too early (~72 hours after first batch of eggs laid) or at the appropriate time (96 hours after egg-laying) and scored if larvae were eaten or accepted. For both treatments, we provided the breeding pair with ten larvae. Following the addition of larvae, we observed the focal individuals for at least 30 minutes to detect acceptance or infanticide of larvae. If the focal individuals did not respond in this time, we checked pairs every 30 minutes to identify the onset of acceptance or infanticide. In some cases, one parent would accept larvae while the other parent committed infanticide; in these cases, we sampled and categorized the male and female separately according to the phenotype that was observed for each. We collected focal individuals while attacking or eating larvae for the Infanticide treatment; individuals in the Accept treatment were collected immediately after the parent made antennal contact with the larvae and a feeding event was observed. This manipulation often led to a discrepancy in the temporal cues used by males versus females, as females base their window of larval acceptance on when eggs were laid (Oldekop et al. 2007), and males likely base their window on timing and frequency of female acceptance of mating. Thus, to reach an adequate sample size, we removed the female after the first batch of eggs were laid, which increased the likelihood that males would accept larvae. We collected n = 20 for both sexes in both treatments.

#### Parental Care

We performed four experimental treatments: Nesting (pair mated and have prepared the mouse “nest”, no larvae), Care (mated, have nest, and are providing care to larvae), Post Care (mated, have nest, and have provided care until larvae disperse), and Double Care (mated, have nest and larvae for two breeding attempts) to test the hypothesis that *IT* or *ITR* influence parental care. We collected n = 20 of each treatment.

We collected whole heads of males and females 24-36 hours after the male had been reunited with the female for the Nesting treatment. For individuals in the Care treatment, we allowed eggs from the second batch to hatch and crawl to the brood ball naturally to ensure parental acceptance of larvae. We checked each breeding box every 12-24 hours to determine when larvae arrived. After parents interacted with their larvae for 24 hours, we scored parental care behavior (see *Behavioral observations*). We collected whole heads of the focal male and female 24 hours after the larvae were first observed on the carcass (at age Day 21-23). Individuals in the Post Care treatment were allowed to complete parenting. Following the dispersal of larvae away from the carcass and the cessation of parental care, we housed the male and female singly in individual containers with soil for 24 hours prior to collection.

To determine whether changes of *IT* and *ITR* expression were directly related to the expression of care, and/or organized the capacity for expression of parental behavior across the lifetime, pairs were allowed to breed twice, creating a Double Care treatment. We removed Double Care beetles for 24 hours following larval dispersal of first breeding attempt, and then put the pair with a new mouse. We allowed the pair to breed normally and monitored the set up every 24 hours to determine egg lay date and the arrival of larvae. 24 hours after hatch, we performed the behavioral observations following the same methods as the Care treatment (see *Behavioral Observations*). We provided a new mate to the focal individual when mortality of one parent occurred during or immediately after first breeding attempt (n = 7 of 20 breeding attempts).

#### Behavioral Observations

We conducted focal behavioral observations to determine how variation in parental care was related to variation in *IT* and *ITR* by observing each pair in the Care and Double Care treatments. We removed lids to breeding boxes 30 minutes prior to observation to allow pairs to acclimate and conducted all observations under red lights, as *N. orbicollis* are nocturnal (Wilson et al. 1984). All observations were by a single observer. We recorded direct care (male or female regurgitated food to larvae) and indirect care (carcass cleaning and maintenance). We also recorded time spent showing “no care”, defined as the parent on the carcass but not engaged in direct or indirect care or was off the carcass. We collapsed these data to align with previous research using scan sampling, where we recorded the activity of each beetle every minute for the 30-minute observation. This gave each individual three scores (direct care, indirect care, and no care) which together summed to 30. Continuous observation allowed for the observer to categorize behavior into direct or indirect care by placing the behavior at each time point into its functional context. Male parental care is variable and often depends on the presence and activity of the female (Walling et al. 2008; Benowitz and Moore 2016). For males that did not show direct parental care during the initial behavioral observation, we removed females to induce male parental care, and then collected males while they were showing direct care. For males that did not show either indirect or direct parental care during the behavioral observations (n = 13), we removed and collected the female to initiate care behavior in the males. Males were checked every 2 – 4 hours to determine when there was a transition to feeding behavior. For these males, we verified direct care of larvae prior to collection by observing at least one larval feeding event.

### Gene Expression Analysis for *Inotocin* and *Inotocin Receptor*

We decapitated the focal beetle with dissecting scissors, flash freezing the whole heads in liquid nitrogen (Roy-Zokan et al. 2015), and immediately placing in storage at −80 °C until RNA extraction. We homogenized heads in liquid nitrogen using a mortar and pestle and then extracted RNA from each sample using Qiagen’s RNAeasy Lipid Tissue Mini-Kit (cat. no. 74106) following (Roy-Zokan et al. 2015), including the DNase I (Qiagen) treatment. We quantified RNA using a Qubit 2.0 Fluorometer (Invitrogen Corporation, Carlsbad, CA, USA) and the RNA Broad Range protocols per manufacturer’s instructions. We synthesized cDNA with Quanta Biosciences qScript reverse transcriptase master mix (Quanta Biosciences, Gaithersburg, MD, USA) following the manufacturer’s instructions from 500 ng total RNA. RNA was stored at −80 °C and cDNA was stored at −20 °C.

We performed quantitative real-time PCR (qRT-PCR) for each of our genes of interest and an endogenous control genes (*gapdh*) following methods of Cunningham et al. (2014) and using transcriptome of Benowitz et al. (2017). We performed qRT-PCR with SYBR I Green Master Mix and a Roche LightCycler 480 (Roche Applied Science, Indianapolis, IN, USA) following the manufacturer’s protocol with triple technical replicates of 10 μl reactions (n = 20 for each parental context, n = 10 for each water treatment and age group, for both sexes). Primers were at a working concentration of at 1.33 μmol/l. We used an annealing temperature of 60° C during the amplification cycles. We established the stability of endogenous reference gene amplicons by visual inspection and analysis of *C*_T_ values found that *gapdh* did not vary substantially across social, water, or age contexts. The Minimum Information for Publication of Quantitative Real-Time PCR Experiments (MIQE) data can be found in Appendix 1.

### Statistical Analyses

All statistical analyses were performed using JMP Pro (v. 16.0.0) and all figures were produced in SigmaPlot (v. 14.5). Results are means ± SE. We used the ΔΔ*C*_T_ method (Livak and Schmittgen 2001) to convert raw expression data to relative expression values. For all analyses, we used males as the comparison group for male analyses, and females as the comparison group for female analyses. We used virgin Day 21 as the comparison group for the age series. For all behavioral states (Accept vs. Infanticide, Nesting, Care, and Post Care, and Care vs. Double Care), we used the behavioral state that occurred earliest within the experiment as the comparison group (Infanticide, Nesting, and Care, respectively). We used High Humidity as the comparison group for the Low vs. High Humidity experiment. Data were visually inspected for outliers. We tested for the effect of Low vs. High Humidity, Accept vs. Infanticide, and Care vs. Double Care using pooled t-tests, and for the effect of age and parental care states using Analysis of Variance (ANOVA). We used splines for non-parametric visualization of how expression changed with age.

We also assessed if there was a direct association between *IT* and *ITR* expression and parental care behaviors observed in behavioral observations. We pooled data from individuals in the Care and Double Care treatments as there was no difference in *IT* and *ITR* expression between treatments (see Results). We analyzed the association between *IT* and *ITR* expression and care behavior using linear regression for individuals that were observed to feed larvae at least once during the targeted behavioral observation without removing the mate (females: n = 34, males: n = 16). We analyzed the association between *IT* and *ITR* expression and indirect parental care using linear regression for individuals that interacted with the brood ball or larvae in any way (females: n = 34, males: n = 21). For one behavioral observation, the pair buried the carcass, and the observation time was thus only 25 minutes; for this reason, all behavioral data are presented as a proportion of the observation time. We excluded a subset of observations (n = 6) because male and female behavior was more indicative of a response to perceived competition or disruption (i.e., frequent stridulation and both individuals off the carcass exploring breeding box).

## Results

### Inotocin and Water Balance

The relative humidity of the two water balance treatments was lower and higher than the average relative humidity experienced by beetles breeding with a carcass (F_2,57_ = 1032.03, P < 0.0001); i.e., the Low Humidity treatments averaged 15.39 ± 1.3 % RH, breeding beetles with a carcass had a RH of 72.99 ± 0.79 %, and beetles in the High Humidity treatment experienced an average humidity of 84.8 ± 1.3 % RH. Males in the Low Humidity treatment had higher *IT* and *ITR* expression than males in the High Humidity treatment (*IT*: t_1,18_ = −2.99, P < 0.008; *ITR*: t_1,18_ = −2.62, P < 0.02; Fig. 2A). Female beetles in Low Humidity treatments had trend towards higher *IT* (t_1,18_ = −1.96, P < 0.07; Fig. 2A), while Low Humidity females expressed higher *ITR* (t_1,18_ = −2.12, P < 0.05; Fig. 2B), than females in the High Humidity treatment.

**Figure 2.**
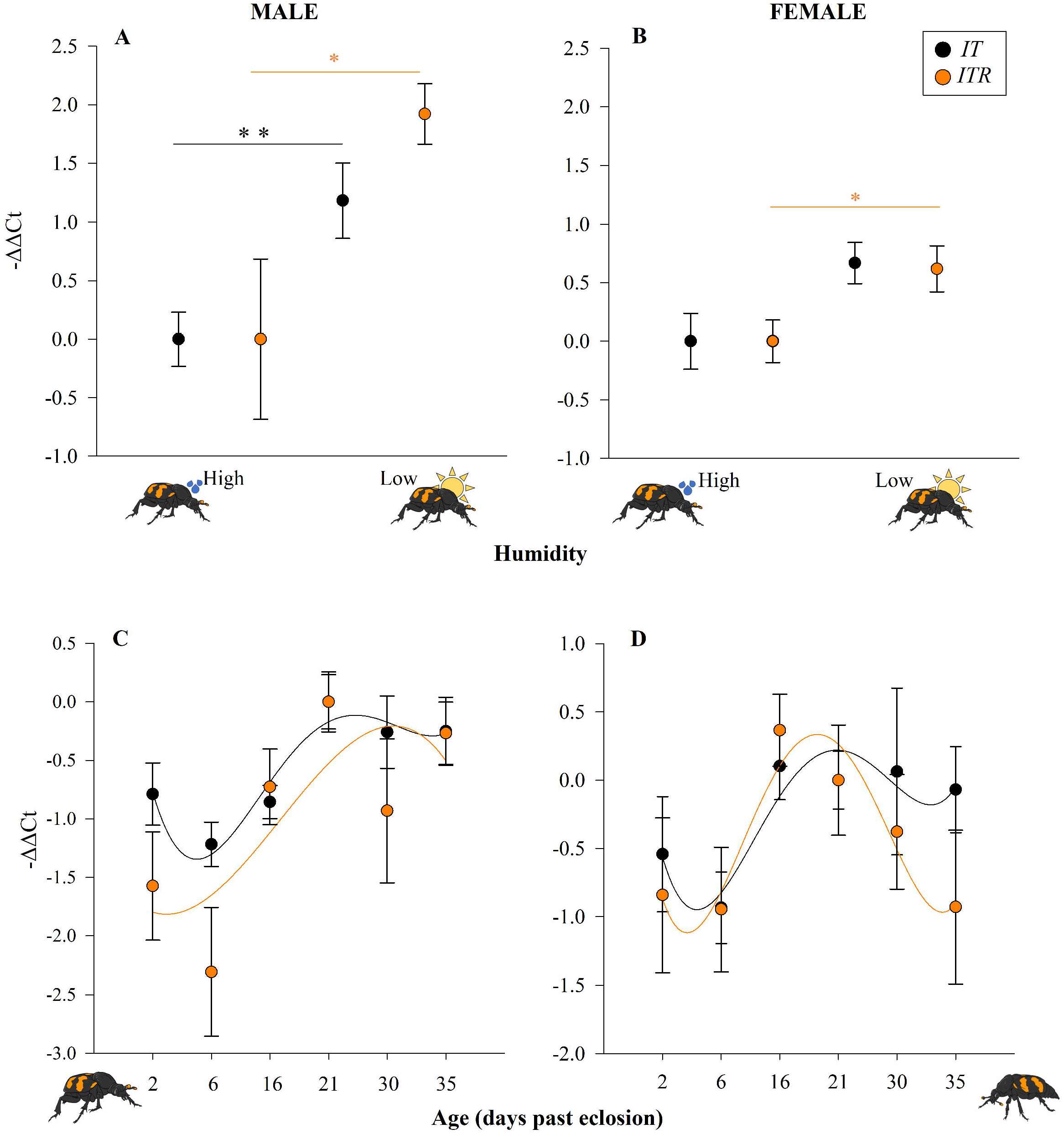
Expression of *IT* and *ITR* genes is associated with water balance and development. Orange circles represent *IT* expression, while closed circles represent *ITR*. (***A***) *IT* and *ITR* were both associated with water balance in males while (***B***) *ITR* but not *IT* was associated with water balance in females. (***C***) Both *IT* and *ITR* change across the lifetime of males, corresponding to the transition to sexual maturity (between Day 6 and 16). Lastly, (***D***) *IT* and *ITR* do not change across the lifetime of females, although there is a trend towards a similar increase as observed in males. For *A & B*, p-values are denoted over the lines, with a P-value less than 0.05 (*), and less than 0.01 (**). For *C & D*, spline curves were used for nonparametric visualization of these patterns. Results are mean ± SEM (n = 10 for each treatment).

### Inotocin in Social Contexts

Both *IT* and *ITR* changed over age for males and females. Splines indicate that both increase around sexual maturity, and plateau at the higher level of expression in males and decrease again at higher ages in females (Figure 2C and D). Males showed increased *IT* (F_5,54_ = 3.45, P = 0.009; Fig. 2C) and *ITR* (F_5,54_ = 3.90, P = 0.004; Fig. 2C) expression across this timespan. In females, neither the expression of *IT* (F_5,54_ = 1.27, P = 0.30; Fig. 2C) nor *ITR* (F_5,54_ = 1.44, P = 0.22; Fig. 2D) changed with adult age, likely because of the decrease at 30 and 35 days of age.

Variation in male parental care was associated with *IT* expression. While males that committed infanticide did not differ in *IT* or *ITR* expression compared to males that accepted larvae (*IT*: t_1,38_ = −0.84, P = 0.41; *ITR*: t_1,38_ = −0.08, P = 0.94), males in the Care treatment had higher *IT* than males in the Nesting and Post Care treatments (F_2,57_ = 4.23, P = 0.02, Fig. 3A; but not *ITR*: F_2,57_ = 0.24, P = 0.79). We found no effect of male parental state on *ITR* (F_2,57_ = 0.24, P = 0.79). Moreover, males that spent more time feeding larvae had higher *IT* expression than males that fed less (F_1,14_ = 8.91, r^2^ = 0.39, P < 0.01; Fig. 3C). There was no relationship between the amount of time males spent feeding larvae and *ITR* (F_1,14_ = 0.08, r^2^ = 0.005, P = 0.79). Amount of indirect care provided was negatively correlated with *IT* expression in males (F_1,19_ = 4.33, r^2^ = 0.19, P = 0.05; Fig. 3C), but there was no relationship between *ITR* and indirect care by males (F_1,19_ = 0.06, r^2^ = 0.003, P = 0.80). Males parenting for the first time did not have higher *IT* or *ITR* than males parenting for a second time (*IT*: t_1,38_ = −0.62, P = 0.54; *ITR*: t_1,39_ = 0.56, P = 0.58).

**Figure 3.**
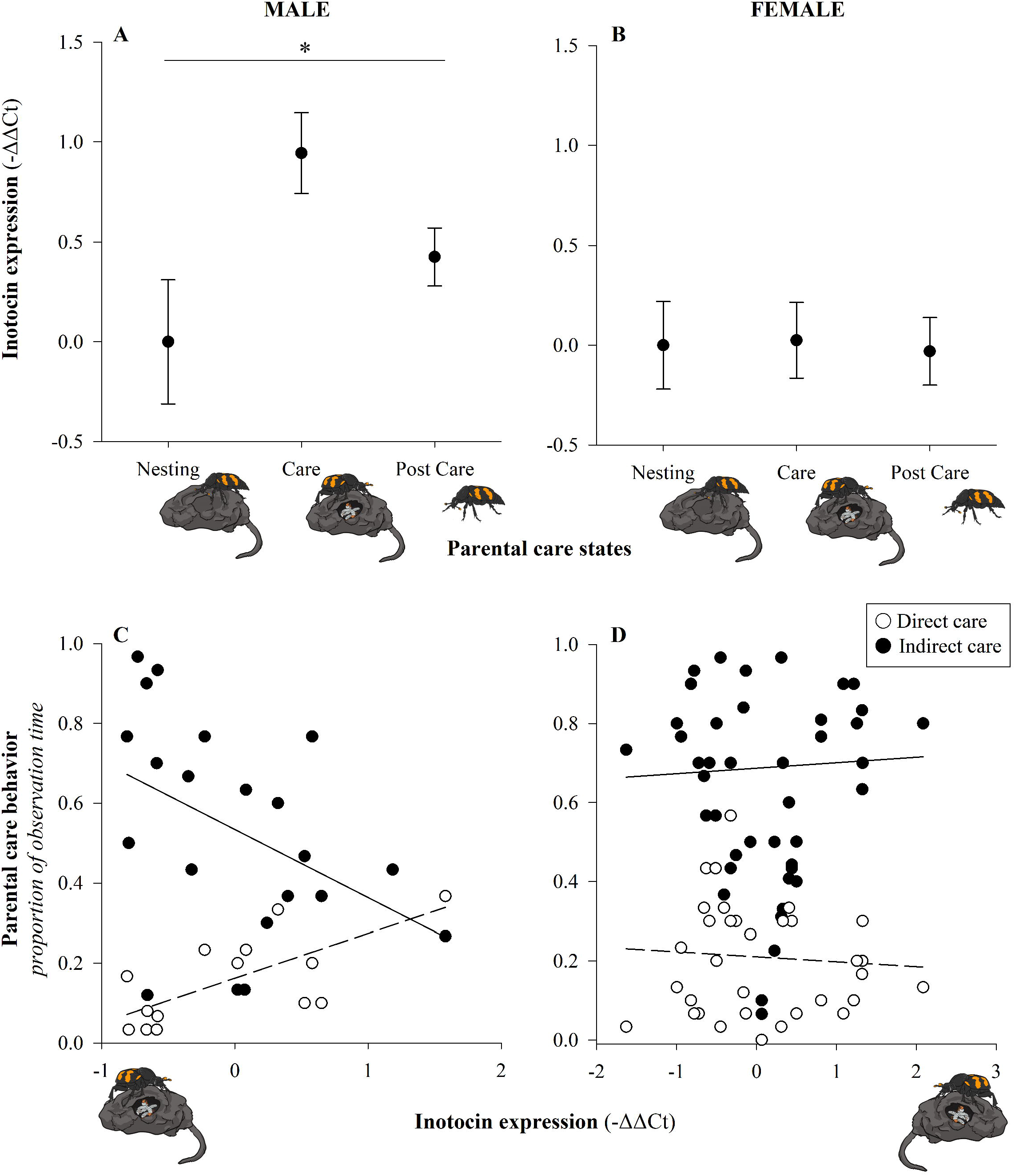
*IT* associated with variation in male but not female parental care. (***A***) Males showed an association between parental care states and *IT* (n = 20 for each state), but (***B***) there was no difference in *IT* for females across parental care states (n = 20 for each). Moreover, (***C***), for the subset of males that performed care behavior without removing the female, males that fed more had higher *IT* (open circles, dashed line; n = 16) and males that spent more time engaged in indirect care had lower *IT* (closed circles, solid line; n = 21). Discrepancy in sample sizes reflects that males that did not show any direct or indirect care were excluded from those respective analyses. On the other hand, (***D***) there was no relationship between *IT* and the proportion of time females spent either feeding larvae (open circles and dashed line) or engaging in indirect care (closed circles and solid line; n = 34). For *A* & *B*, significant P-values are denoted over the lines, with a p-value less than 0.05 denoted with a *. Results are mean ± SEM.

In contrast to males, we found no significant difference in gene expression corresponding to female parenting. Females that committed infanticide did not differ in *IT* or *ITR* expression relative to females that accepted and cared for larvae (*IT*: t_1,38_ = 1.49, P = 0.14; *ITR*: t_1,38_ = 0.34, P = 0.74). Female parental care states were not associated with *IT* (F_2,57_ = 0.02, P = 0.98; Fig. 3B) or *ITR* expression (F_2,57_ = 0.21, P = 0.81). There was no relationship between the proportion of time females spent feeding larvae and *IT* or *ITR* expression (*IT*: F_1,32_ = 0.17, r^2^ = 0.005, P = 0.69, Fig. 3D; *ITR*: F_1,32_ = 3.15, r^2^ = 0.09, P = 0.09). There was no relationship between indirect care provided by females and either *IT* (F_1,32_ = 0.11, r^2^ = 0.003, P = 0.74, Fig. 3D) or *ITR* (F_1,33_ = 0.0004, r^2^ < 0. 0001, P = 0.98). Females parenting for the first time did not have higher *IT* or *ITR* than females parenting for a second time (*IT*: t_1,38_ = 0.37, P = 0.71; *ITR*: t_1,37_ = −0.14, P = 0.89).

## Discussion

Our results demonstrate that inotocin system in *N. orbicollis* is associated with both non-social and social contexts important for the oxytocin/vasopressin system more broadly. As expected, we found that *IT* is associated with water balance in both sexes (Fig. 2), supporting other research indicating that water balance was an ancestral function of the oxytocin/vasopressin system (Fig. 1; Fujino et al. 1999; Ukena et al. 2008; Koto et al. 2019). We further found that *IT* is associated with parenting, like other homologues in the oxytocin/vasopressin system (Fig. 1), but that the associations differ between males and females.

We found that *IT* expression changes as beetles transition to sexual maturity. While the pattern was found in both sexes, it was more pronounced in males than in females (Fig. 2). Pre-reproductive males had lower *IT* expression than post-sexual maturity males, where it plateaued. There was a similar increase in female *IT* expression around sexual maturity. Across vertebrate species, it has been repeatedly demonstrated that oxytocin/vasopressin systems develop in a way that influences subsequent expression of reproductive and parenting behaviors (e.g., Hammock 2015; Vaidyanathan and Hammock 2017). Based on conserved functions observed across vertebrates, it is possible that *IT* is involved in priming beetles for reproduction, but further research is needed to determine how the development of inotocinergic system is related to reproduction.

Rather than a general role in parental care, we found that increased *IT* expression was associated with variation in male but not female parenting in *N. orbicollis*. This holds for both transition from nesting to offspring care and back to a non-parenting state, as well as overall amount of time attending to offspring and performing indirect care activities such as carcass maintenance (Fig. 3). Overall willingness to accept or commit infanticide of larvae was not associated with differences in *IT* or *ITR* expression in either sex, suggesting that *IT* has a role limited to interactions with larvae during direct care. It is possible that *IT* also has a role in female parental care that is obscured because female care is less variable than the care provided by males. One way to resolve this problem is to determine where *IT* is expressed in the brains of females and males, as research in other systems has found that variation in sex-specificity of neuropeptides can be enabled by variation in where the gene is expressed. Previous work has indicated that *IT* is often produced in the SEZ, and *IT* in this location is likely to be related to water balance pathways (Aikins et al. 2008; Fetter-Pruneda et al. 2021). However, it is unknown where *IT* is expressed in the brains of parenting insects, and future research should determine whether *IT* is expressed in similar areas of the brain in both females and males in different care states.

There are three potential explanations for our results: first, that a role in male parenting arose de-novo in burying beetles. This is possible but not parsimonious given the distribution of social influences of oxytocin/vasopressin homologues across vertebrate taxa (Fig. 1). Second, given that both male and female parental behaviors are influenced by the oxytocin/vasopressin system in vertebrates, it could be that the female role of *IT* was lost in burying beetles. However, this is unlikely, given that other studies of female insects have not found a definitive influence of *IT* on parenting behavior. Although changes in *IT* have been documented across a reproductive cycle and putatively associated with parenting (Nagy et al. 2021) and other social behaviors (Fetter-Pruneda et al. 2021), there has been no other experimental manipulation of parental care state relative to *IT* or *ITR*. Finally, it may be that influences on social behavior are ancestral, but we only see an association between male care and *IT* because *IT* is associated with flexibility in social behavior. Male parental care is much more variable than female care in *N. orbicollis*. Genes of the oxytocin/vasopressin family have a complex relationship with social behavior and likely involve multiple genes that perform functionally redundant roles (Berendzen et al. 2022). Yet another explanation is that *IT* shows similar evolutionary lability in the sex-specificity of social effects as oxytocin/vasopressin do in vertebrate species (Fig. 1), although large-scale functional assays across invertebrate species are required to test this.

There is evidence across taxa that support the hypothesis that the oxytocin/vasopressin system influences flexibility in behavior. In *Caenorhabditis elegans*, the single copy homologue for oxytocin/vasopressin, *nematocin*, influences male mating (Garrison et al. 2012) and gustatory associative learning overall (Garrison et al. 2012; Scott et al. 2017). Oxytocin/vasopressin have been found to increase the salience of social signals and group formation in a context-dependent manner in both humans and fish (Kosfeld et al. 2005; Reddon et al. 2012), and are associated with plasticity in socially-mediated behaviors in reef fish (e.g., Godwin et al. 2000). *IT* may perform a similar role and acts to organize parental care behavior in *N. orbicollis* by increasing the relevance of offspring cues to males. Support for this idea is provided by our result that males with higher *IT* fed their larvae more (i.e., increased attention to larvae) than males with lower *IT*, and males with lower *IT* spent more time engaged in indirect care, where attention was focused on the maintenance of the carcass instead of larvae. Thus, the influence of *IT* on male parental care may be more apparent because male parental provisioning is more responsive to environmental context than female care (Smiseth et al. 2005; Moss and Moore 2021). If *IT* is involved in mediating flexibility of parenting, this may explain why a widespread role for IT and parenting has not been found, as many insects with parental care often show little flexibility in the performance of care. For example, while workers transition from feeding and caring for larvae to foraging in eusocial insects, they often do so in a temporal fashion moving from one task to the next in a developmental series. This limits the scope for determining how variation in behavior is associated with variation in *IT* expression.

Our results also raise questions about the role of gene duplication in behavioral evolution. The vertebrate nonapeptides oxytocin and vasopressin evolved from a tandem duplication event of a vasopressin-like gene (Van Kesteren et al. 1995; Goodson 2008; Theofanopoulou et al. 2021) that occurred approximately 500 million years ago in the common ancestor of vertebrates (Yamashita and Kitano 2013). While the role of this system in parenting has been demonstrated across a wide diversity of vertebrate taxa (Fig. 1, SI Table 1), both peptides demonstrate great evolutionarily lability in which function is associated with males and females (Donaldson and Young 2008; Goodson and Thompson 2010; Kelly and Goodson 2014; Kohl et al. 2018). The breadth of functions associated with oxytocin/vasopressin and their sex-specificity was thought to be a result of gene duplication, as gene duplication is thought to facilitate the evolution of new gene functions. *IT* is a single copy homologue and yet also shows a sex-specific association with parenting, which may indicate that the sex-specificity of these neuropeptides, or their social functions, do not require gene duplication.

Multiple lines of evidence support the idea that the ancestral gene was more similar to *vasopressin* (Van Kesteren et al. 1995; Goodson 2008) and *IT* is more structurally similar to the *vasopressin* group than *oxytocin* (Stafflinger et al. 2008). Oxytocin was first discovered in female vertebrates in association with parturition (Dale 1909), and correspondingly, there has been a greater emphasis on the role of oxytocin in females with a lesser emphasis on vasopressin and males. For this reason, the discovery that *IT* is associated with variation in male parental care behavior, but not female parental care, was unexpected. Nevertheless, the results of our study support the idea that the social influences and potentially the sex-specificity of this system are likely ancestral.

## Acknowledgements

We thank Kathryn Kollars for producing the beetle art. We thank Hans Otto, Kyle Benowitz, Shannon Harris, Emily Shelby, and Kathryn Kollars for discussions and/or feedback on the manuscript. A.L.P was funded by a USDA cooperative agreement to AJM and A.T.B. was funded by a Natural Sciences and Engineering Research Council of Canada discovery grant to Marla Sokolowski.

## Author Contributions

A.L.P. and A.J.M. conceived and designed the experiments. A.L.P. conducted experiments. A.L.P., E.C.M., P.J.M., A.T.B., and C.B.C. conducted laboratory work. A.L.P. and A.J.M analyzed the data. All authors contributed to the writing of the manuscript.

## Data Availability

The data underlying this article are available in the Dryad Digital Repository (https://doi.org/10.5061/dryad.q573n5tn4).

