## Supplementary Table 1 for "Insect homolog of oxytocin/vasopressin associated with parenting of males but not females in a subsocial beetle"

|  |  | Oxytocin | | Vasopressin | | One copy | |
| --- | --- | --- | --- | --- | --- | --- | --- |
| Function | Taxa | Female | Male | Female | Male | Female | Male |
| Affiliation | Birds | (1) | (2, 3) | (4-6) | (4-7) |  |  |
|  | Fish | (8-10) | (8, 11-14) | (10) | (11, 12) |  |  |
|  | Mammals | (15-19) | (14, 15, 20, 21) | (22) | (21, 23, 24) |  |  |
| Aggression | Birds | (25) | (26) | (25, 27) | (28, 29) |  |  |
|  | Fish | (30) | (30) | (31) | (31-33) |  |  |
|  | Mammals | (16, 34) | (35) | (15, 34, 36) | (15, 23, 36, 37) |  |  |
| Nest building | Birds | (38) | (38) | (38, 39) | (38, 39) |  |  |
|  | Invertebrates | - | - | - | - | (40) |  |
|  | Mammals | (41) | (42) | (41) | (41) |  |  |
| Myoactivity | Birds | (43) | - | (43, 44) | - |  |  |
|  | Fish | (45) | (46) | (47, 48) | (46) |  |  |
|  | Invertebrates | - | - | - | - | (40, 49) |  |
|  | Mammals | (50) | (51) | (52) | (51) |  |  |
| Direct parental care | Amphibian | (53) | (54) | (54) | (54) |  |  |
|  | Fish | (55) | (33, 56) | (57) | (33, 54, 57, 58) |  |  |
|  | Invertebrate | - | - | - | - | (59) |  |
|  | Mammals | (41, 60, 61) | (14, 21, 62) | (41, 60, 63) | (21, 64, 65) |  |  |
| Water balance | Birds | (66) | (67) | (66, 67) | (68) |  |  |
|  | Fish | (69) | (69) | (69-71) | (69-71) |  |  |
|  | Invertebrates | - | - | - | - | (72, 73) | (72, 73) |
|  | Mammals | (74) | (75) | (74) | (75) |  |  |

**Supplementary Table 2.** **Oxytocin/vasopressin homologs often perform similar functions in both sexes across species**; however, each often have sex-specific functions within species. Studies for which oxytocin and/or vasopressin homologs have been found to be associated with each function, including changes in gene expression, peptide abundance, and/or receptor distribution. This is not an exhaustive review, and discrepancies between the taxa represented (e.g., birds are not represented under “direct parental care”) generally reflects a lack of studies that explicitly test the function of each neuropeptide in both males and females. Accordingly, reference was given to taxa for which the function of each was tested for both sexes.
