## Supplementary Figure 1 for "Insect homolog of oxytocin/vasopressin associated with parenting of males but not females in a subsocial beetle"

**Supplementary Information Figure 1.** Experimental design.
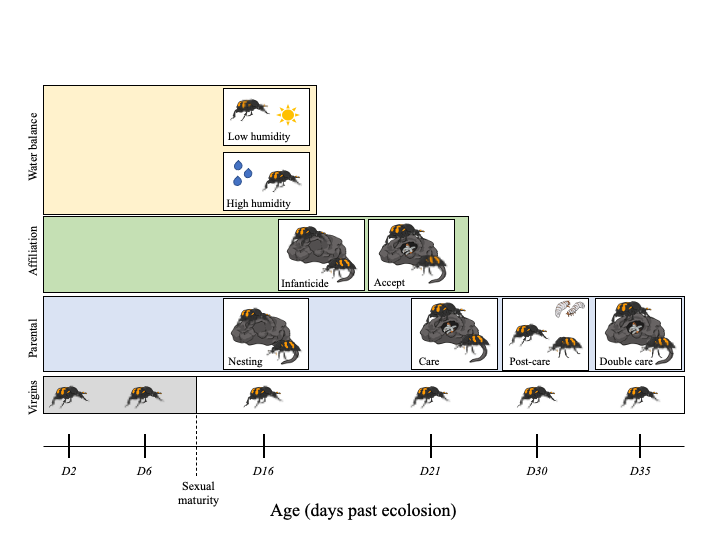

**SI Figure 1.** Experimental design. Isolated, virgin adult male and female beetles were collected for an age series (Days 2, 6, 16, 21, 30, and 35 past eclosion). For parental states, beetles were collected at Nesting (mated, eggs, carcass), Care (peak larval care 24 hours after hatch), Post Care (larvae dispersed, parents isolated for 24 hours after larval dispersal), and Double Care (peak larval care 24 hours after hatch for a second breeding). For Affiliation, beetles were induced to accept/feed larvae (Accept), or to attack/eat larvae (Infanticide) by manipulating temporal cues. Lastly, water balance was manipulated by exposing beetles to either a Low or High Humidity treatment.
