## Supplementary Appendix 1 for "Insect homolog of oxytocin/vasopressin associated with parenting of males but not females in a subsocial beetle"

**Supplementary Information Appendix I**

Additional information about the quantitative real-time PCR protocol

I. Additional information suggested in the Minimum Information for Publication of Quantitative Real-Time PCR Experiments (MIQE) guidelines not already provided in this paper.
A. Quantitative real-time PCR (qRT-PCR) primer sequences

*inotocin* - NCBI accession GGAA01017632.1

1. forward: ACTCTTGCCTCATCACCAAC
2. reverse: CGAGATGCACGGTTTGATTTG

*inotocin receptor* – NCBI accession GGAA01004254.1

1. forward: GCAACTTTGGCCATCATCTTC
2. reverse: AGTTTCTTCCGACCTGCATAC

*gapdh* – NCBI accession GGAA01002248.1

1. forward: CGACTACATGGTGTACCTGTTC
2. reverse: TGCCGTTGACGATGAGTTT

*tbp –* NCBI accession KY654102.1

1. forward: TATGGTGGGCAGTTGTGATG

2. reverse: ACGGTAAATCAAACCAGGGAATA

Primers were manufactured by Integrated DNA Technology (IDT, Coralvill, IA, USA) and purified with IDT's standard desalting technique. We used the software IDT PrimerQuest^TM^ Tool for development of qRT primers.

A. qRT-PCR validation

Primer efficiency:

*inotocin:* E=2.04, r^2^=0.9355

*inotocin receptor:* E=2.05, r^2^=0.9382

*tbp:* E=2.13, r^2^=0.9533

*gapdh:* E=1.95, r^2^=0.9996
